# Fast and interpretable quantification of biological shape heterogeneity via stratified Wasserstein kernel

**DOI:** 10.1101/2025.11.05.686865

**Authors:** Wenjun Zhao, Danica J. Sutherland, Khanh Dao Duc

## Abstract

Modern imaging technologies produce vast collections of cellular and subcellular structures, calling for principled methods that enable shape comparison across individuals and populations. We introduce the stratified Wasserstein framework, which treats each shape as an unstructured point cloud and embeds it into Euclidean space via ranked local distance profiles. This embedding yields an isometry-invariant Euclidean distance and a positive-definite kernel for population analysis, with a consistent sample-based estimator that supports large datasets in near-quadratic time. By leveraging kernel methods, the framework enables statistically rigorous tasks such as nonparametric hypothesis testing, providing theoretical guarantees as well as interpretability. We demonstrate our framework’s applicability to large-scale biological datasets. Analyzing 2D cancer cell contours, we quantify population-level discrepancies and identify representative cells contributing most strongly to the observed differences. Using 3D volumes of cell envelope and nucleus, we reveal progression patterns that capture morphological changes across cell populations both at the level of individual shapes. These results establish a simple and principled tool for population-level biological shape analysis, with potential impact across diverse domains of computational imaging and data science.

## Introduction

Recent advances in high-throughput imaging have generated massive collections of biological shapes across multiple scales, from whole tissues to single cells, subcellular structures, and protein conformations. These datasets enable population-level analyses of morphology across conditions, but they also expose a methodological gap: we lack tools that are both interpretable and computationally efficient for comparing shapes at the level of individual objects and at the level of distributions of objects. Questions such as how morphology evolves through the cell cycle [49] or how protein structures differ across species [11] call for comparisons that respect intrinsic geometry, scale to large datasets, and connect naturally to statistical analysis and machine learning.

Traditional approaches for shape comparison face well-known limitations. Many methods rely on reducing data to specific shape features, such as volumes, heights, persistent barcodes in topological data analysis [30], spherical harmonics [22], with features of interest being established *a priori*. Landmark-based analysis pipelines quantify the pairwise discrepancy through the correspondence of specific landmark points, which are manually placed points and can be labor-intensive, subjective, and difficult to reproduce [14] for large-scale datasets. Other classical parametric methods are tailored to particular domains [12, 46], and/or require heavy pre-processing including down-sampling, alignment, or interpolation [23, 47]. In contemporary imaging pipelines with thousands to millions of shapes, these constraints become bottlenecks.

While most existing work has also focused on pairwise distances or shape alignment at the individual level, population-level metrics between shape ensembles remain underexplored. Historically, this gap can be attributed to the relatively small size of shape datasets, where analysis was limited to tens or hundreds of samples. With modern imaging pipelines now producing thousands to millions of shapes, the need for scalable and statistically principled population-level distances has become pressing. A few population-level approaches have been proposed. For example, Fréchet mean and distances to all shapes [29], linear subspace embeddings and Kullback– Leibler divergence [4] have been considered in specific contexts, but they are tied to specific assumptions, and none provide a general-purpose, theoretically grounded, and computationally efficient framework for comparing populations of arbitrary shapes.

In this context, optimal transport theory provides a principled way to compare shape objects given as unstructured point cloud or their probability histograms. In particular, Gromov–Wasserstein distances compare shapes through their internal pairwise distances and avoid explicit alignment [31]. Related lines of work summarize shapes by distributions of distances, either globally or around each point [6, 17, 31, 32]. However, it is well-known that such computations can be intractable [25] and expensive [42], and consequently, they are hard to be applied to datasets at scale. Moreover, it is difficult to build positive definite kernels directly from these distances in dimensions greater than one, which limits downstream statistical tools such as kernel hypothesis tests [20, 21, 51] and representation learning [38, 44]. Existing works [24, 36] reconcile the issue by slicing the measure and reducing the transportation problem to one-dimensional, however, that requires sampling angles of slicing and may not fully utilize the information from the whole shape.

We address these challenges by introducing the *stratified Wasserstein distance*, a simple yet highly effective framework that embeds each unstructured point cloud into an Euclidean space. We show that the distance is Hermitian and produces a kernel that is positive definite and, under mild regularity conditions, characteristic. The construction is invariant to isometries by design and is injective up to isometry. Computationally, the method is nearly quadratic in number of points per shape and outperforms the Gromov–Wasserstein, which is cubic after acceleration via entropic regularization. We illustrate the utility of the framework across diverse biological settings on 2D and 3D shape datasets, focusing on both individual and population-level shape analysis.

## Results

We begin by presenting an overview of the stratified Wasserstein framework, describing its construction and key properties. This provides the conceptual and theoretical foundation for the remainder of the paper. We then demonstrate its performance on two and three-dimensional biological imaging datasets, highlighting its ability to handle diverse shape types and to support population-level statistical analyses such as hypothesis testing and dependence detection.

### Overview of the stratified Wasserstein framework

We propose stratified Wasserstein, a framework that embeds into Euclidean space each shape, representd as an unstructured point cloud, and facilitates kernel methods in that space for population-level quantification tasks. Compared to existing shape distances such as Gromov–Wasserstein and its lower bounds [31] as defined in equations M1, M2, and M3 (Methods), the proposed framework achieves similar discriminative power while being computationally more efficient, with complexity nearly quadratic in the number of points and empirical runtimes typically below 1% of those required by Gromov–Wasserstein methods. The induced kernel is characteristic, so population-level statistics via kernel methods retain their standard statistical guarantees, including consistency and power against alternatives [20, 21, 45]. Figure 1 provides a schematic overview of our procedure, that takes local distance distribution to produce shape embeddings and derive population-level statistics. Here we describe the main construction and summarize its key properties; detailed statements and proofs are deferred to Section and the Supplementary Information.

**Figure 1.**
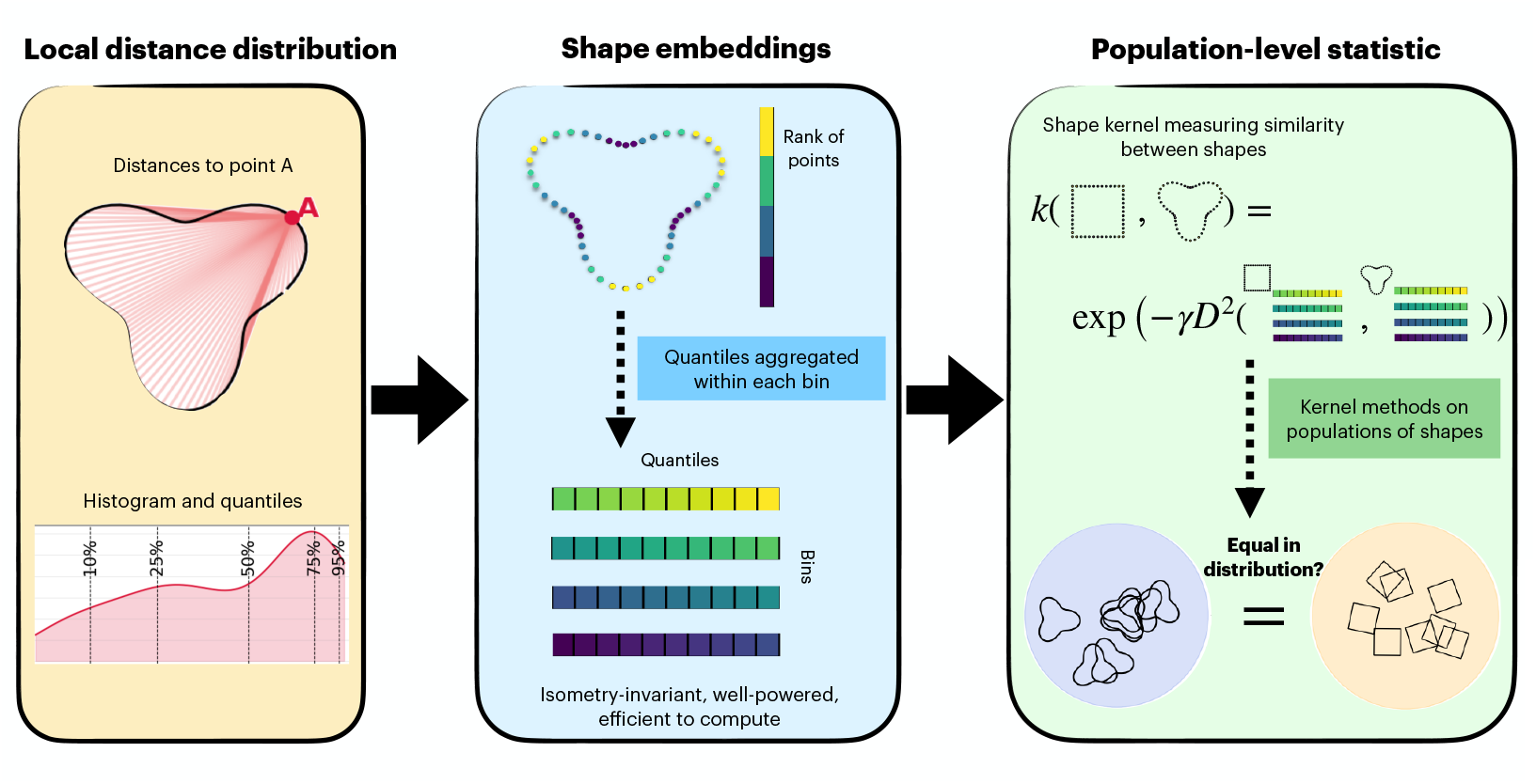
A summary of our procedure. Given unstructured point cloud in 2D or 3D representing a shape of a wide range of types, we first compute the intrinsic distances on each shape and localized for each point. Using these local distances, a wide range of downstream tasks can be performed given a large population of shapes, such as dimension reduction, clustering, hypothesis testing, and feature selection. The detailed methodology is presented in subsequent sections, with rigorous theoretical foundation discussed.

First, we represent each shape by a point cloud together with an intrinsic distance, such as the geodesic distance on the surface or the Euclidean distance in ambient space. For every point on the shape, we compute the distribution of distances to all other points, which captures its local geometric structure. We then rank the points according to a continuous functional of these local distance distributions, which induces a one-dimensional stratification of the shape. Within each stratum, we summarize the local geometry by recording a set of quantiles of the distance distributions. This defines an embedding of the shape into a two-dimensional function space indexed by the stratification variable and the quantile level. The stratified Wasserstein distance between two shapes is simply the *L*^*p*^ distance between their embeddings, computed over this two-dimensional domain.

In this work, we show that the stratified Wasserstein distance inherits many of the desirable properties of Gromov–Wasserstein while significantly improving computational efficiency. It is naturally invariant to isometric transformations because of the invariance property of local distance distributions. Under mild regularity conditions on the ranking functional used for disintegrating the shape, the distance is injective up to isometry: two shapes have zero distance if and only if they are isometric. Because the distance is induced by an *L*^*p*^ norm in the embedded space, standard kernels such as the Gaussian and Laplace built from it are positive definite. When the distance is injective, these kernels are also characteristic, meaning they can distinguish any two distributions of shapes. The estimator of the distance based on finite point clouds is statistically consistent: as the number of points per shape grows, the estimated distance converges to the population distance. The computational cost scales nearly quadratically with the number of points per shape, which is far more efficient than Gromov–Wasserstein methods.

Finally, because the kernel is characteristic, standard population-level statistics built from it inherit strong theoretical guarantees. For example, maximum mean discrepancy (MMD) [20] measures any form of discrepancy between two populations, while the Hilbert–Schmidt Independence Criterion (HSIC) [21] measures the strength of dependence between variables. Both statistics are consistent and have power against all fixed alternatives, which enables rigorous nonparametric two-sample testing and dependence detection between shape distributions and external covariates.

We provide Table 1 to compare our proposed stratified distance between shapes against other relative alternatives. Compared to the other distances, stratified Wasserstein is Hilbertian, can be consistently estimated from finite samples, and achieves the lowest asymptotic computational complexity. It is conditionally injective and yields a characteristic kernel if the oracle functional for sorting and binning is chosen carefully. The balance between discriminative power and computational complexity is further demonstrated in numerical examples on real world biological shape datasets as discussed in the next section. Empirical comparison on these distances can be found in Supplementary Figures S3 and S4.

**Table 1:**
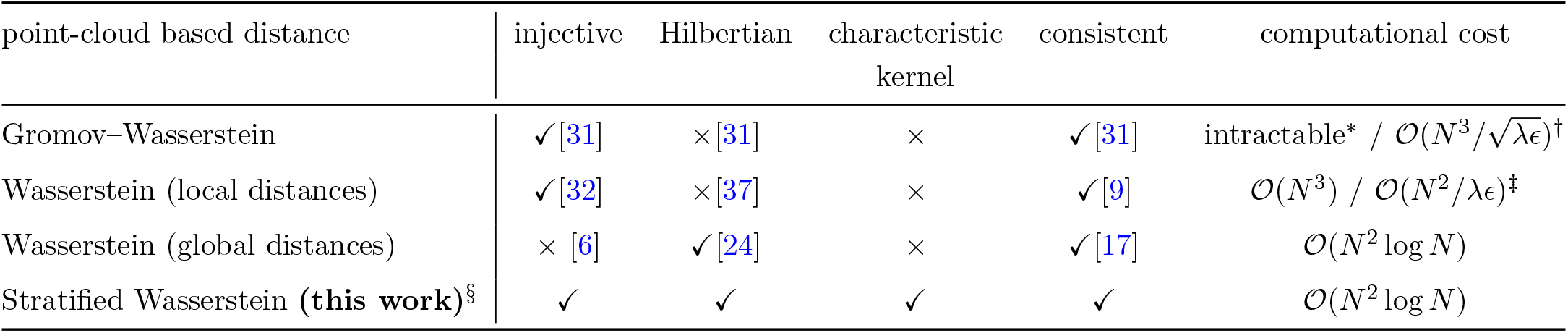
Comparison between distances with shapes represented as point clouds in 2D/3D. For Gromov–Wasserstein and Wasserstein between local distances, computational cost for both exact computation (left) and entropic approximation (right) are both shown. N = points per shape. ^*^Exact GW is NP-hard [25]; ^†^Entropic GW complexity depends on λ (regularization) and ϵ (tolerance) [41], both are small, positive values for accurate approximation; ^‡^Systematic studies on computational costs can be found in [3, 10, 13]. ^§^ Our framework has a conditionally injective and characteristic kernel under Hypotheses (H1) and (H2) with details in Appendices.

### Breast cancer cell contour shapes from fluorescence microscopy

We apply our framework to 2D cancer cell shapes obtained from fluorescence microscopy [1, 2]. Cell images are binarized, and their boundaries are extracted to form discrete curves given by the 2D coordinates of the cell contours. The dataset includes cancer cell shapes from three different cell lines for breast cancer, corresponding to: (1) MCF10A (228 cells): non-tumorigenic human breast mammary gland epithelial cell line, which is a classic model for normal cells, (2) MCF7 (225 cells): breast cancer line with relatively low metastatic potential, and (3) MDA-MB-231 (abbreviated MDA, 224 cells): a triple-negative breast cancer cell line that is highly invasive and commonly used as a model for metastatic progression. Given populations of these cell contours, the goal is to test whether the underlying shape distributions differ significantly across groups defined by cell lines. Understanding these shape differences may help reveal whether and how cell morphology encodes functional behaviors for different cancer types. An example of the raw images, as well as random samples of cells from each cell line are visualized in Figure 2, panels (a) and (b), and more images can be found in Supplementary Figure S5.

**Figure 2.**
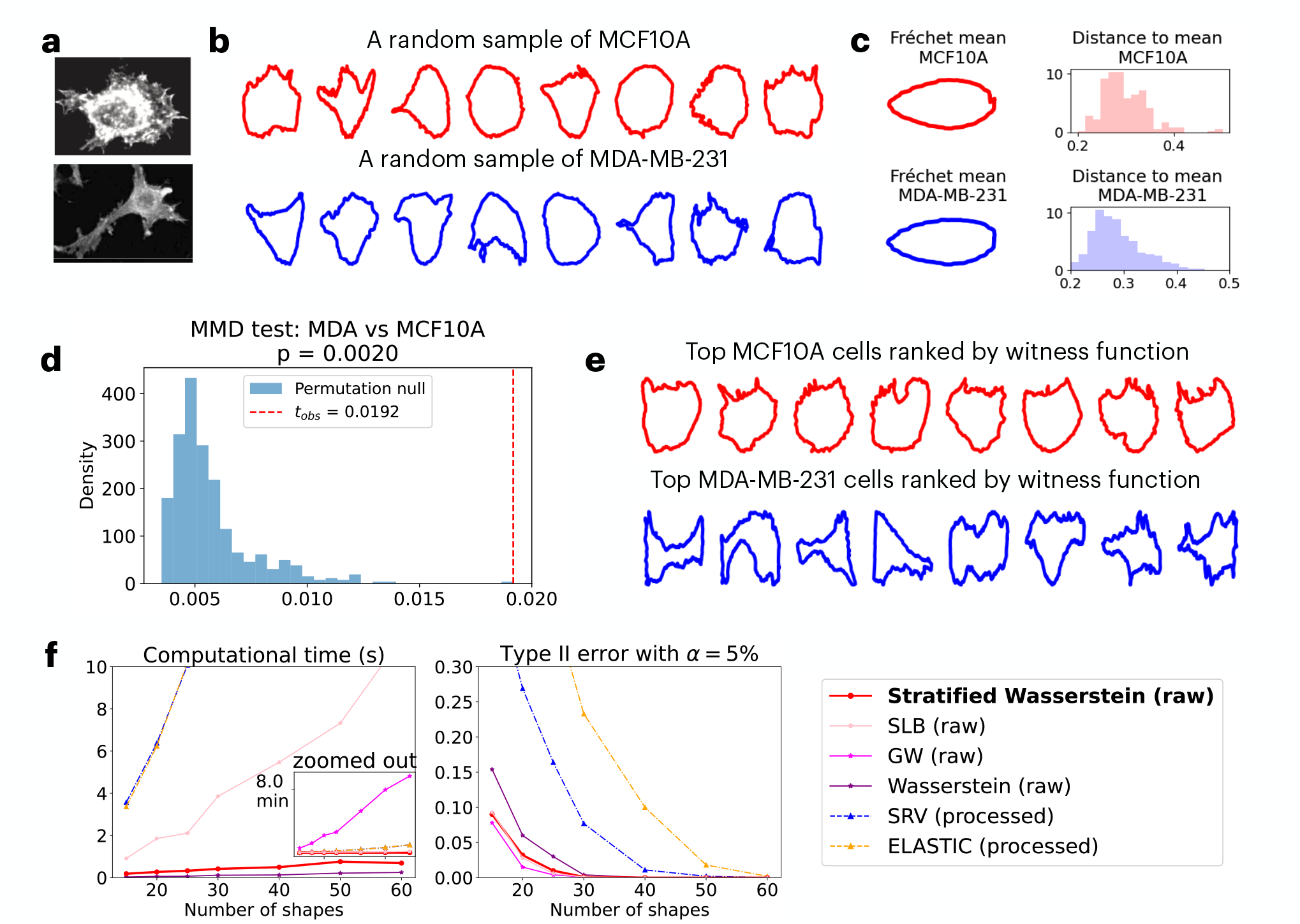
Case study on 2D cancer cell shape populations represented by contours. (a) Example fluorescence microscopy images of cell lines curated by [1]. (b) Cell contours after segmentation, with a random sample of 8 shapes from each population visualized. (c) Inspection of first and second moments suggests similarity between the MCF10A and MDA populations, as shown via Fréchet mean shape and distances to the mean shape within each population under the SRV metric, computed using Geomstats [29]. (d) To further probe differences between MCF10A and MDA, we compute the MMD-based test statistic and its p-value using our proposed stratified kernel, leveraging all available cell shapes. (e) Cells with the top 8 witness scores in each population highlight those whose shapes differ most from the respective population majority. (f) Empirical probability of Type II error and computational time under small (15) to moderate (60) sample sizes, benchmarked across various kernel-based tests.

### Relative MMD test reveals similarity between triple negative cancer cells and non-cancerous cells

Upon testing relative MMD between three populations, we observed that: Despite being a cancer line, MDA resembles MCF10A (normal cells) more than MCF7 (low-metastasis cancer). The empirical estimate of squared MMD reveals that MMD^2^(MCF10A, MDA) = 0.0189, much smaller than the estimate of MMD^2^(MCF7, MDA) = 0.1409. To test whether the difference is statistically significant, we perform a relative MMD test for the following:

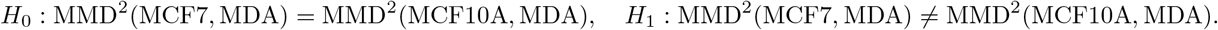

The *p*-value is computed using a permutation test that pools MCF7 and MCF10A, under the null hypothesis that the two populations are equally distant from MDA. Specifically, we permute the MCF7/MCF10A labels and recompute the difference in squared MMD to MDA for each shuffle. With 1000 permutations, this test results in a *p*-value of 0.001 (Supplementary Figure S5), suggesting that the MDA line (high metastatic cancer) is morphologically closer to MCF10A (normal cell line) than to MCF7 (low metastatic cancer). This observation is consistent to findings from gene expression studies [34], which reported that both MCF10A and MDA share a basal-like subtype, while MCF7 belongs to the luminal class. Our shape-based results suggest that cell morphology is more closely aligned with molecular subtype (basal versus luminal) than with cancer status.

The result is also consistent with the prior approach [26] that computes Fréchet mean shape under the square root velocity (SRV) metric [46] through geomstats [29] package. The mean shapes of processed contours from all 3 populations are shown in Panel (b) of Supplementary Figure S5, where MCF10A and MDA mean shapes are both elongated and almost coincide.

### Absolute MMD test indicates discrepancy between populations on the tails

Given almost identical mean shapes and distribution of distance to mean shown in Panel (c) of Figure 2, it seems that the two shape populations of MDA and MCF10A agree up to mean and variance. A natural follow-up question is whether the generating distributions of the two shape populations are truly identical. To this end, we zoom in on the unexpected pair and perform an MMD test on those two populations, which detects arbitrary order of discrepancy in distribution that extends beyond second order:

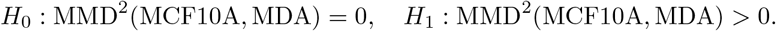

With 1,000 permutations, it shows a *p*-value of 0.002, suggesting a significant difference in the two populations. To localize the morphological differences, we examine extreme shapes identified by the MMD witness function, which highlights regions where the shape distributions differ most. Top 8 cells in each population with the highest values of the MMD witness function are visualized in Figure 2, panel (e). The extremal subset of MDA cells exhibits highly irregular and protrusive morphologies, which are key contributors to the shape-based distinction from MCF10A. These shapes are likely enriched in invasive behaviors as known for MDA [18]. Notably, the statistical difference in MMD is not driven by the mean shape or the distances to mean shape (Figure 2, panel (c)), but by this tail population in MDA, which can only be captured using nonparametric test like the kernel MMD.

### The tests are computationally efficient and statistically well-powered

To assess the power of the absolute and relative tests, we perform randomized experiments to quantify the empirical Type I and II error rates. We compare our kernels to the framework suggested by Zhang et al [52] using square-root velocity (SRV) metric and its variant (Elastic) for MMD. The benchmark approach is performed on preprocessed data, where each shape has been interpolated to have 2,000 points and aligned to the same reference shape beforehand. In contrast, in our approaches, we perform the test directly on raw data without any processing.

We perform the tests for sample sizes varying between 15 and 60 via MMD. To systematically quantify the error, for each sample size, we resample and repeat the experiments 1,000 times, compute the *p*-value via permutation test, and report the relative frequency of making an error in Figure 2 (panel (f)). For Type II error, we draw one sample from each group, perform the test, and compute the relative frequency that the equality of shape distributions cannot be rejected. Among our proposed distances, Gromov–Wasserstein consistently achieves the lowest Type II error, followed by stratified Wasserstein distances and the second lower bound (SLB) of Gromov–Wasserstein. All Wasserstein-type distances outperform SRV-based metrics at sample sizes under 40. Similar experiments and observation on error rates for both Type I and II are done for relative test and shown in Supplementary Figure S5. We also note that the error rate is robust to the choices of hyperparameters, and sensitivity results are reported in Supplementary Figure S6.

To assess computational efficiency, we record the wall time required to compute all pairwise distances between shapes in each subsample. These results are shown in Figure 2 (f). While GW offers the best statistical power, its computational cost is substantial. Our stratified Wasserstein achieves a favorable balance: while offering second best Type II error control, slightly worse than Gromov–Wasserstein, it is significantly more efficient than Gromov–Wasserstein and its second lower bound, and is second only to the pooled Wasserstein method, which simply aggregates and sorts all pairwise distances.

In sum, our nonparametric, distribution-level approach operates directly on raw contour shapes, detects tail effects, and localizes the drivers of discrepancy while remaining computationally practical.

### Allen Institute 3D Cell and Nucleus Shapes

In this example, we demonstrate that quantiles of distance matrices encode meaningful morphological information, using 3D cell imaging data from Viana et al. [49]. We analyze a subset of 5,764 cells used by the original authors to train a classifier, including annotations for cell types reflecting six mitotic stages (M0, M1M2, M3, M4M5, M6M7 early, and M6M7 half) as well as three outlier types (blob, dead, wrong). Existing dimension reduction methods rely either on hand-crafted features (e.g., volume and height of the cell and nucleus) [49] or on learned latent representations from deep models such as variational autoencoders [8], followed by UMAP. However, these methods are either highly specialized for the application or require significant training effort.

### Dimension reduction for shapes reveals relevant morphological variations

We pre-process each binary image by eroding it to extract surface points, followed by down-sampling to retain 200 points per cell. For each cell, we divide it into 100 bins, compute 100 quantiles for both the cell shape and the nucleus shape in each bin, concatenate them, and apply UMAP to the resulting 20,000-dimensional vector. Figure 3 panel (a) shows 2D UMAP embeddings obtained using stratified Wasserstein distance. To benchmark our result, we performed dimension reduction using 4 additional methods and visualized them in panel (b) of Figure 3 that are based on (1) Euclidean distance in raw binary images, (2) dominant features through volumes of cell and nucleus, (3) the intermediate latent layer using a pre-trained PointNet model [39], and (4) Gromov-Wasserstein distance with each cell downsampled to 100 points due to prohibitive computational cost. Our dimension reduction best preserves the cyclic and continuous nature of the shape dynamics, consistent with the feature-based and PointNet embeddings, but with a more smooth trajectory that reflects the remark on the ambiguity of manual annotation by Viana et al. [49].

**Figure 3.**
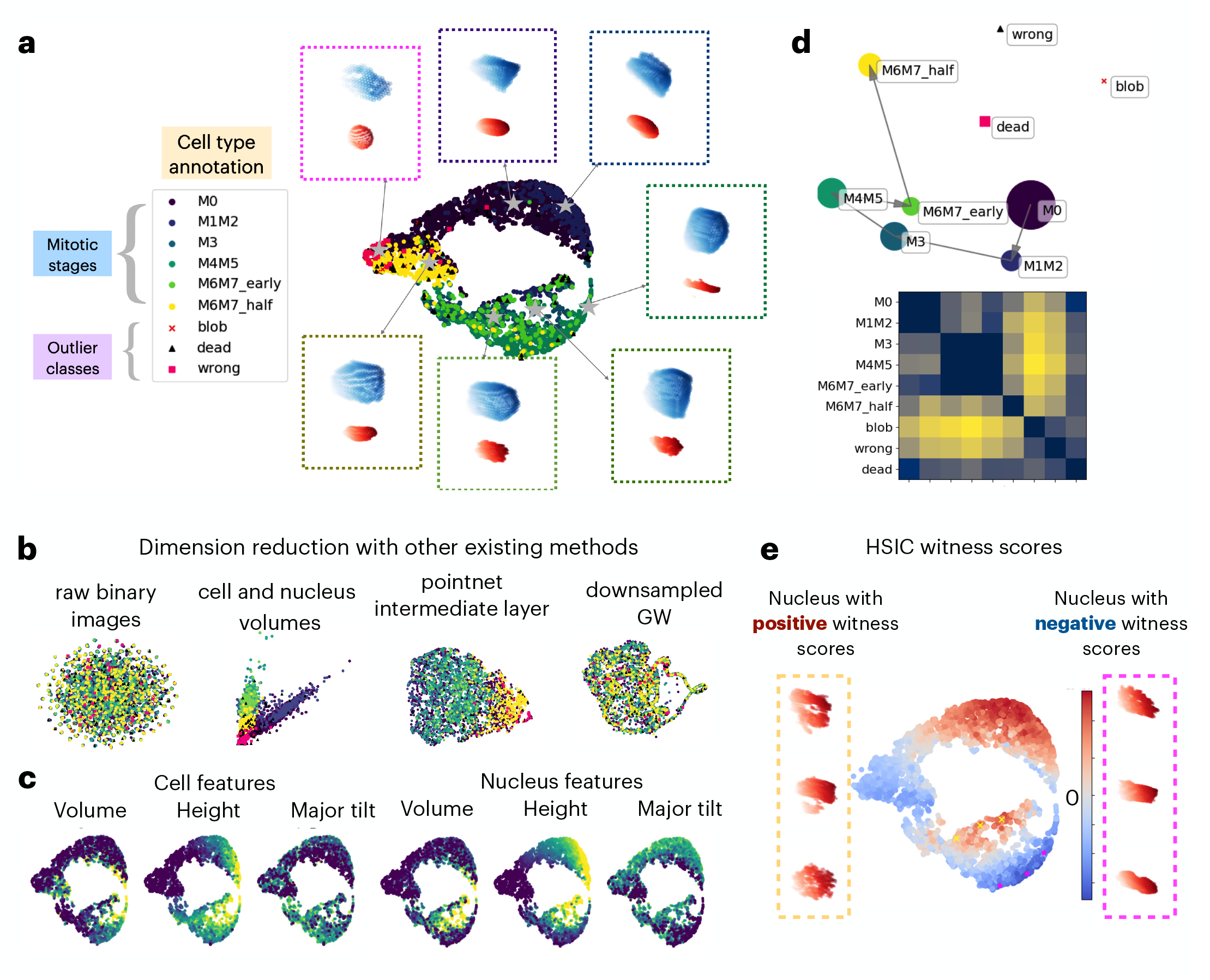
Case study on 3D cell and nucleus shape populations. (a) UMAP embedding of the annotated mitotic dataset from Viana et al. [49] through stratified local distance distributions, and representative cell (blue) and nucleus (red) shapes along the cycle. (b) Cell-level embeddings from standard methods fail to reveal dynamics. Euclidean distances between raw binary images and Gromov–Wasserstein distances on downsampled point clouds show no meaningful pattern, while feature-based embeddings (volumes or neural networks) capture heterogeneity but not the cyclic continuum of mitosis. (c) Cell features displayed against UMAP allows for interpretation of shape changes within each stage. (d) Population-level embedding using proposed MMD reveals cyclic process consistent to (a). (e) HSIC witness scores for individual cells, revealing regions with mixed positive and negative values near late mitotic stages. We selected 3 cells with high positive witness scores (yellow crosses) and 3 with high negative witness scores (magenta stars), and display the nucleus shapes on the sides.

This embedding provides biological insight into morphological changes during the mitotic cycle. In Figure 3(c), we color cells by morphological features to show how these evolve along the cycle. Starting from the M1M2 phase, both cell and nucleus volumes increase significantly, followed by a marked rise in nucleus height during M3 and M4M5. These trends align with the known stages of mitosis, where volume increase reflects DNA and organelle replication, and structural transitions (e.g., chromatin condensation and nuclear envelope breakdown) are hallmarks of the later phases [50]. The outlier cells have particularly smaller volumes and heights in both cell and nucleus shapes, likely reflecting incomplete mitosis, abnormal states, or segmentation artifacts.

### Embeddings of populations respects mitotic cycle progression

To better summarize discrepancies between shape populations corresponding to different mitotic stages, we embedded the cell populations using MMD as a distance metric between groups. Figure 3, panel (d) displays the resulting distance matrix, which exhibits a banded structure: populations from adjacent stages (e.g., M0 and M1M2) have lower distances, while outlier cell types are farther from the mitotic populations. Since MMD defines a valid population-level distance, we further applied multidimensional scaling (MDS) to embed the populations in 2D space, revealing a clear progression through mitotic stages. We note that two alternative population metrics, energy distance (and Wasserstein distance), do not perform as nicely as MMD-based embedding, likely due to the fact that the kernel methods are more resistant to noise through smoothing, which is crucial for real imaging data. Comparison to other candidates of population-level distances can be found in Supplementary Figure S7.

### HSIC witness scores locate representative shapes for mitotic progression

HSIC witness scores, much like the witness function used in MMD-based testing, provide a way to interpret how individual samples contribute to the overall dependence between variables. We exclude the outlier cell types, and used continuous values of mitotic stages to better capture the periodic nature by assigning equally spaced values from 0 to 2*π*, with von Mises kernel *k*^*γ*,periodic^(*y*_1_, *y*_2_) = exp(*γ* cos(*y*_1_ − *y*_2_)). When applied to cell and nuclear shapes across the cell cycle, these scores highlight specific regions where the shape is particularly informative or uninformative of mitotic stage (Figure 3, panel (e)). To better understand these regions, we visualized representative cells with the strongest positive and negative witness scores. Interestingly, although all selected cells belong to late mitotic stages, their nuclear morphologies differ substantially. Cells with positive witness scores (yellow crosses) tend to show bi-lobed or segmented nuclear shapes, consistent with active division, while those with negative scores (magenta stars) exhibit smooth, undivided nuclei. This suggests that the HSIC witness score captures subtle morphological differences that reflect how well shape aligns with mitotic progression, helping to identify stage-aware prototypes with distinct developmental states.

Overall, our framework recovers the known mitotic trajectory while pinpointing rare, stage-specific morphologies, demonstrating the potential to reveal unknown heterogeneity within data.

## Discussion

To provide tools for quantifying heterogeneity in shape data at both the individual and population levels, we proposed a stratified Wasserstein framework to embed shape data into a Euclidean space and utilize kernel methods therein. The construction is naturally invariant under isometric transformations and admits a consistent sample estimate for both shape–shape distances and between-population distances via maximum mean discrepancy (MMD). On 2D breast-cell contours and 3D mitotic cell and nuclear segmentations, the framework supports dimensionality reduction, clustering, and nonparametric hypothesis testing, matching or surpassing state-of-the-art methods while substantially reducing runtime. Although we focus on smooth planar curves and surfaces, the formulation extends to other metric–measure data types, e.g., neuron trees modeled as metric graphs, when equipped with a geodesic distance and mild regularity [19, 32]. Because it operates directly on unstructured point clouds with variable sizes and requires no landmarks or global alignment, the approach offers a unified and scalable route to shape analysis across domains. More broadly, our framework has potential applications across domains that involve the quantification of shape/graph data, such as brain images, protein density maps, and social networks.

Next, we outline several limitations and potential avenues for improvement. First, in practice we rank points by the mean of their local distance distributions; on near-symmetric geometries this statistic can be weakly discriminative. While the performance is comparable to Gromov–Wasserstein and second lower bound for random polygons (Supplementary Figure S3), the shape distance under the stratified framework is significantly overestimated on T-shapes with symmetry (Supplementary Figure S4). A principled remedy is a lexicographic rank using the first *k* moments (or quantiles) of the local distance laws; we show such rankings become injective on finite samples as *k* → ∞, but selecting a small, data-adaptive *k* and a corresponding ranking functional remains open. Second, the numbers of bins and quantiles are user-chosen without specific guidelines. Deriving finite-sample error bounds for the stratified estimator would enable data-driven defaults that balance discretization bias against bin variance. Third, the kernel bandwidth is currently set by the median-distance heuristic or ad hoc tuning; under low signal-to-noise, a common pitfall in current imaging, more principled rules (e.g., maximizing estimated MMD test power or kernel alignment, or noise-aware plug-ins) could improve performance. Addressing these issues would further strengthen robustness and ease of use for practitioners.

The proposed framework has the potential to facilitate new computational tools and biological discoveries in multiple ways. By connecting shape space to kernels, the framework unlocks a broad toolbox for inference and representation. (i) With multimodal measurements (e.g., shapes and transcriptomics), kernel-based conditional independence (KCI) [51] can yield *p*-values for identifying driver genes that explain morphology while controlling confounders, aiding inference on gene regulation underlying shape change. (ii) Coupled with kernel representation learning [38] and functional data analysis (e.g., kernel PCA or Gaussian processes on the embedding), one can learn low-dimensional surrogates that capture discrete phenotypes or continuous trajectories along which shapes vary most rapidly. (iii) Given its computational efficiency, the framework can be integrated into in vivo perturbation screens to prioritize conditions that induce the largest distributional shifts in shape. As a whole, these analyses turn the framework into a practical engine for hypothesis testing, causal inference, perturbation discovery, and more–at scale.

By addressing the challenges listed above and further extending the framework to other tasks, stratified Wasserstein has the potential to become an even more powerful tool, enabling more comprehensive insights into modeling the highly heterogeneous shape space across diverse biological contexts.

## Methods

### Shape representation and intrinsic distances

We represent each shape as an unstructured point cloud 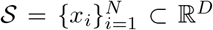 with *D* ∈ {2, 3}. Each shape is equipped with an intrinsic distance *d*_*S*_ : 𝒮 × 𝒮 → ℝ_≥0_ and a uniform measure *µ*_*S*_. Distances can be chosen flexibly, depending on the application, including Euclidean distance, geodesic distance on a *k*-nearest neighbor graph [19], or diffusion distance [28]. The intrinsic distance matrix is *C*_*ij*_ = *d*_*S*_(*x*_*i*_, *x*_*j*_). For scale invariance, shapes are rescaled so that the median pairwise distance equals one.

### Gromov–Wasserstein and related distances

#### Gromov–Wasserstein distance

For two metric measure spaces (S_*i*_, *d*_*i*_, *µ*_*i*_), *i* = 1, 2, the *p*–Gromov–Wasserstein (GW) distance [7, 31] is defined by

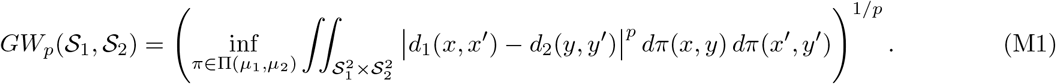

In the discrete setting, this corresponds to minimizing over coupling matrices between points. Exact computation is NP-hard [25], and entropic regularization is typically used to obtain approximate solutions with complexity 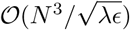 [41]. In all numerical experiments, we used the implementation provided by Python Optimal Transport [15, 16].

#### Global Wasserstein distance

A simpler lower bound compares the global distributions of pairwise distances [6, 17, 35]. Flatten the distance matrices of two shapes into vectors *c*_1_, *c*_2_, and let 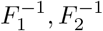 be the quantile functions of their empirical distributions. The *p*–Wasserstein distance between these distributions is

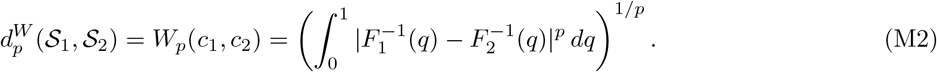

This distance can be computed efficiently in 𝒪(*N* ^2^ log *N*), but it is not injective: different shapes may yield the same global distance distribution.

#### Second lower bound (SLB)

A stronger metric compares local distance distributions [31, 32]:

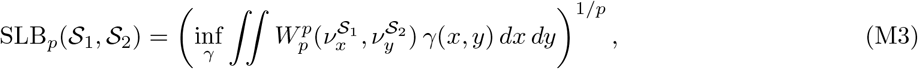

where 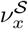 is the local distance distribution at point *x* and *γ* is a coupling between points. SLB is strictly stronger than the global Wasserstein distance and is injective for smooth closed shapes under regularity assumptions [32], but remains computationally demanding and is not Hermitian.

### Stratified Wasserstein distance

The proposed stratified Wasserstein distance combines the geometric discriminative power of GW-type distances with the efficiency of quantile-based embeddings. For each point *x* ∈ 𝒮, we compute its local distance distribution 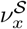. A continuous ranking functional *r* _𝒮_ (*x*) (e.g., the mean of 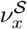) induces a stratification of the shape via its normalized rank *U* ^𝒮^ (*x*) = *F* (*r* _𝒮_ (*x*)), where *F* is the cumulative distribution function of *r*_𝒮_. The shape measure *µ*_𝒮_ is disintegrated along *U*^𝒮^,

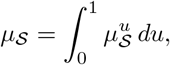

where 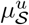 denotes the conditional law at stratum *u*. We define the function *Q* by evaluating the quantile function of *ν* at level *q* ∈ [0, 1]:

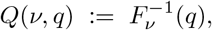

where 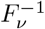 denotes the (left-continuous) generalized inverse of *F*_*ν*_. For each stratum *u* ∈ [0, 1] and quantile level *q* ∈ [0, 1], we compute

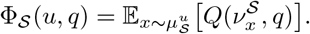

Overall, we have defined an embedding Φ_𝒮_ ∈ *L*^2^([0, 1]^2^). The stratified Wasserstein distance between two shapes is the *L*^*p*^ distance between their embeddings:

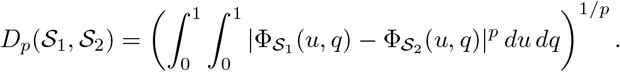

Under mild regularity conditions on the ranking functional, the distance is injective up to isometry (Theorem 2). Since it is induced by an *L*^2^ norm, Gaussian and Laplace kernels constructed from *D*_2_ are positive definite, and if the embedding is injective, these kernels are characteristic. The empirical estimator of the distance, based on binning and quantiles, is statistically consistent as the number of points grows [17]. Its computational complexity is nearly quadratic in the number of points, offering substantial savings over GW and SLB.

### Benchmarks used for comparison

#### Elastic metrics between 2D planar curves

In the task on 2D cancer contour shapes, for comparison, we used the elastic metric with two choices of parameters [5, 27, 33]. Let *γ* : [0, 1] → ℝ^*d*^ be a smooth parameterized curve, and *h*_1_, *h*_2_ be two tangent vector fields along *γ*. The *elastic metric* with parameters *a, b >* 0 is defined by

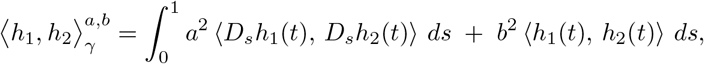

where *s* denotes the arc-length parameter of *γ*, and

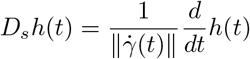

is the derivative of *h* with respect to arc-length. The square root velocity has *a* = 1*/*2 and *b* = 1, while the default elastic metric implemented by Geomstats [29] has default *a* = 1 and *b* = 1*/*2.

#### Pointnet distance between 3D images

As a modern deep learning-based benchmark, we used the feature encoding layers of a pointnet++ model [39, 40] that was pretrained on ModelNet40 classification task, accessed at https://guochengqian.github.io/PointNeXt/modelzoo/. Image of each cell and nucleus is first down-sampled to 2,048 points, and embedded into a feature space of dimension 1,024. We use this embedding for dimension reduction and Euclidean distance therein for MMD as benchmarks of our 3D cell image case study.

#### Features of 3D cell shapes

We computed features of 3D cell shapes for benchmarking and visualization purpose, specifically, for volume, height, and major tilt. Each image is centered at the centroid and applied principal component analysis to determine its dominant axes of variation. The volume is defined as the vertical extent (difference between maximum and minimum) along the third coordinate that has the minimum variance. The volume is computed for the convex hull of the points. The major tilt is computed by the angle between the dominant principal axis and the third, vertical axis.

### Kernel methods on shape populations

Positive-definite kernels constructed from the stratified Wasserstein distance enable population-level statistical analysis using kernel methods. We focus on maximum mean discrepancy (MMD) and Hilbert–Schmidt independence criterion (HSIC), which are widely used nonparametric statistics with strong theoretical guarantees.

#### Maximum mean discrepancy (MMD)

MMD [20] is the squared distance between kernel mean embeddings of two distributions *P*_*A*_ and *P*_*B*_ in the reproducing kernel Hilbert space (RKHS) associated with kernel *k*:

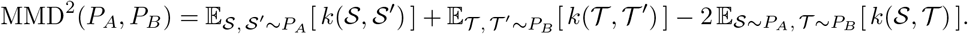

Given samples 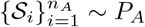 and 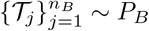, the (biased) empirical estimator is

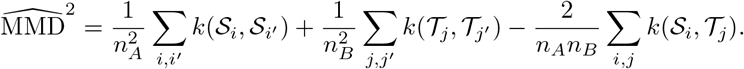

Significance is assessed by permutation testing of group labels. When *k* is characteristic (as is the case for Gaussian or Laplace kernels on the stratified Wasserstein metric), MMD equals zero if and only if *P*_*A*_ = *P*_*B*_. The test is consistent against all fixed alternatives, meaning its power converges to one as sample sizes grow [20, 48].

#### MMD witness function

The MMD witness function [20]

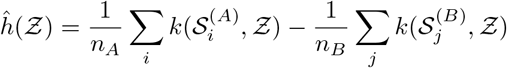

identifies regions of shape space where the two distributions differ most strongly. Evaluating 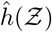 at sample shapes highlights representative shapes that contribute to the observed differences.

#### Hilbert–Schmidt independence criterion (HSIC)

HSIC [21] quantifies dependence between random variables using kernels on each domain, and can be interpreted as the maximum mean discrepancy (MMD) between the joint distribution *P*_𝒮,*Y*_ and the product of its marginals *P*_*S*_ ⊗ *P*_*Y*_. For paired data 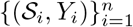, with char-acteristic kernels *k* on shapes and *ℓ* on covariates, HSIC is defined as the squared Hilbert–Schmidt norm of the cross-covariance operator in the associated RKHSs:

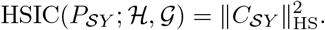

With Gram matrices *K*_*ij*_ = *k*(𝒮_*i*_, 𝒮_*j*_), *L*_*ij*_ = *ℓ*(*Y*_*i*_, *Y*_*j*_), and centering matrix 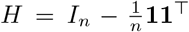, the (biased) estimator is

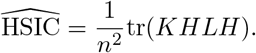

HSIC is zero if and only if 𝒮 and *Y* are independent, provided both kernels are characteristic. It is consistent against all alternatives and detects arbitrary nonlinear dependencies [21, 44].

#### HSIC witness function

Similar to MMD, HSIC admits a witness function that localizes the contribution of individual shape–covariate pairs to the overall dependence. Given kernels *k* and *ℓ* centered in RKHS, the empirical HSIC witness is:

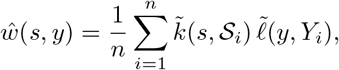

where 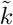 and 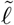 are the centered kernels. Positive values indicate that the pair (*s, y*) supports the observed dependence, while negative values indicate opposition. It is precisely the MMD witness function applied to the joint distribution and the product of marginals. This function can be used to interpret the contributions of specific shapes or covariate values.

### Statistical analyses

All population-level comparisons were performed using kernel methods derived from the stratified Wasserstein distance. For each dataset, Gaussian kernels were constructed using the median heuristic for bandwidth selection on pairwise stratified Wasserstein distances.

Two-sample tests between shape populations were carried out using the maximum mean discrepancy (MMD) statistic [20]. For each test, we used a permutation test with 1,000 permutations to estimate the null distribution, and reported p-values based on the proportion of permuted statistics exceeding the observed value. The MMD witness function was used to localize representative shapes contributing to significant differences between groups.

To assess dependence between shape distributions and external covariates (e.g., developmental stage), we used the Hilbert–Schmidt independence criterion (HSIC) [21] with Laplace kernels on both shape and covariate domains, with bandwidth chosen via the median distance.

For clarity, all the tests in this section are performed using Laplace kernels between shapes with median bandwidth heuristic: 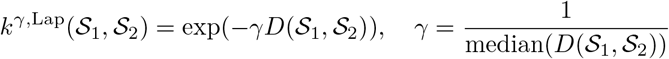, median taken over all possible pairs of shapes, where *D*(𝒮_1_, 𝒮_2_) represents a distance in shape space, such as Gromov–Wasserstein and its alternatives.

For the 3D mitotic cell shape data, the mitotic stage variable consists of six ordered categories (<monospace>M0</monospace>, <monospace>M1M2</monospace>, <monospace>M3</monospace>, <monospace>M4M5</monospace>, <monospace>M6M7 _early</monospace>, <monospace>M6M7 _half</monospace>), which were encoded as integers 0, …, 5. The mitotic stage label was encoded as an angular variable *θ* = 2*π* stage*/*6, and mapped to circular coordinates [cos(*θ*), sin(*θ*)] on the unit circle. A Gaussian kernel applied to these circular embeddings,

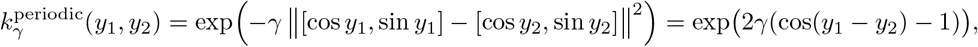

is equivalent (up to a constant factor) to a von Mises kernel (under a rescaling of *γ*)

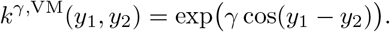

The bandwidth parameter *γ* was selected using the median heuristic, that is, *γ* was set to the median of squared pairwise distances between stage embeddings on the unit circle. This encoding preserves the cyclic topology of the mitotic progression while allowing the stage variable to interact naturally with Euclidean distances between shapes.

Permutation testing over 1,000 trials of the covariate labels was used to assess significance. The HSIC witness function was used to visualize shape–covariate pairs with the strongest contribution to the dependence.

## Data availability

This study did not generate new data. Both processed (aligned and resampled) 2D cancer cell shape data are available at https://github.com/wxli0/dyn/tree/main%4092c7a58/dyn/datasets/breast_cancer. The 3D mitotic cell shape dataset is available at https://open.quiltdata.com/b/allencell/packages/aics/mitotic_annotation.

## Code availability

A repository containing Python code for the 2D and 3D cell shapes for reproducing the results is available at https://github.com/WenjunZHAOwO/StratifiedShapes.

## Appendix

A Description and properties of stratified Wasserstein distance

**Definition 1.1 (Local distance distribution)** *For each shape* 𝒮 *and point x* ∈ *S, the local distance distribution is denoted as:*

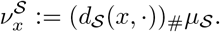

It describes the distribution of all distances *d*(*x, y*) from the reference point *x* to any *y* ∈ 𝒮, and is supported in a bounded interval [0, diam(𝒮)]. From the theory of moment-generating functions, each 1-D distribution is fully determined from all of its moments *m*^*k*^(*x*) for *k* = 1, 2, …, and the existence of moments is guaranteed from the compact support. Moreover, the local distances have no atoms, from the fact that all shapes are continuous.

First, we assume that an oracle ranking function *r* that labels all points is known a priori. This is reasonable, for instance, given that the local distance distributions 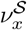 are uniquely determined by their moment sequences 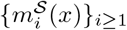, so a functional that achieves lexicographical ordering according to moments would satisfy the criteria:

### Hypothesis (H1) (Ranking functional)

*There exists a measurable functional ψ* : 𝒫 ([0, *diam*(𝒮)]) → ℝ *such that* 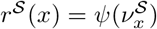 *and:*

1. ( Continuity*) ψ is continuous at* 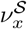 *for µ*_*S*_*-a*.*e. x with respect to weak (equivalently W*_1_*) convergence on* 𝒫 *([0, diam(S)]);*
2. ( A.e. injectivity*) r*^𝒮^ *is injective µ*_*S*_*-a*.*e*.;
3. ( Non-atomic pushforward*) r*^𝒮^#*µ*_*S*_ *has no atoms, so U* = *F* (*r*) *is uniformly distributed on* [0, 1];
4. ( Empirical stability*) For the empirical law* 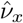 *built from distances*, 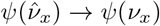 *a*.*s. for µ*_*S*_*-a*.*e. x (hence* 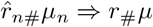).

Such an oracle ranking function does not exist in practice, yet it is always possible to construct one when finite points are available for each shape. Below we propose an empirical surrogate which can be constructed based on finite points and moments of their local distance distributions:

#### Theorem 1

**(Ranking by moments)** *For any shape* S, *the local distance distributions* 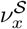 *are uniquely determined by their moment sequences* 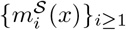. *Moreover, for any finite set of points* {*x*_*a*_} *and for almost every ϵ >* 0, *the polynomial*

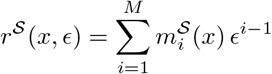

*is injective on* {*x*_*a*_} *whenever the corresponding distributions* 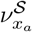 *are distinct*.

**Proof**. Moment determinacy on compact intervals ensures that different 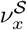 differ in some finite prefix of moments. For such a pair (*x, y*) the difference *r*^𝒮^ (*x, ϵ*) − *r*^𝒮^ (*y, ϵ*) is a nonzero polynomial with finitely many roots. Excluding these exceptional *ϵ*’s, the map *r*^𝒮^ (·, *ϵ*) is injective on the finite sample.

**Remark 1.1:** When *ϵ* is small, this is similar to hierarchical sorting given the moments according to its order. First, we sort all points in S by *m*_1_(*x*), mean distance to all other points. In case of tie, say, there exists a set of points that have equal *m*_1_(*x*), the points are further sorted according to *m*_2_(*x*). Recursively, this will determine an order of all points on the shape, unless there are identical local distance distributions (and it is not necessary to determine the order between them any more).

**Remark 1.2:** The remark assumes no asymmetry. In real data with noise, to distinguish finitely many shapes, each with finitely many points, it often suffices to take *M* = 1 and *ϵ* = 0, so the ranking function only looks at the average distance to all other points.

Using the ranking functional, we can stratify the shape and construct an embedding from it into the Euclidean space, described below.

#### Definition 1.2

**(Stratified Wasserstein distance)** *Let U*^𝒮^ (*x*) = *F* (*r*^𝒮^ (*x, ϵ*)) *be the normalized rank, uniformly distributed over* [0, 1] *within each shape. We first disintegrate each shape measure along U*^𝒮^:

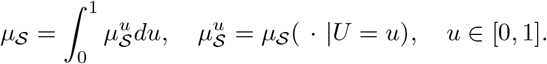

*Then we embed each shape using:*

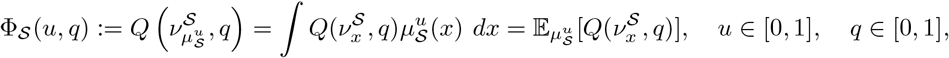

*where Q*(·, *q*) *is the q-th quantile of the local distance distribution. For any shape* 𝒮, *the embeddings* Φ_𝒮_ ∈ *L*^2^([0, 1]^2^) *from the boundedness of* 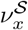. *The stratified distance is defined as the Euclidean distance in the embed-ding space:*

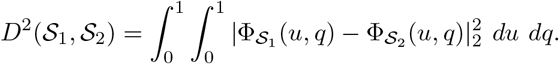

#### Theorem 2

**(Injectivity)** *The stratified distance D*^2^(S_1_, S_2_) *equals zero if and only if* S_1_ *and* S_2_ *are identical up to isometry*.

**Proof**. (⇒) If 𝒮_1_ and 𝒮_2_ are isometric, their local distance distributions coincide, hence the embeddings Φ coincide pointwise, so *D*^2^(𝒮_1_, 𝒮_2_) = 0.

(⇐) Suppose *D*^2^(𝒮_1_, 𝒮_2_) = 0. Define the rank–preserving coupling

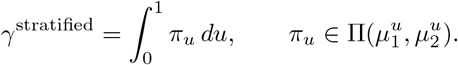

Following Hypothesis (H1)(ii)–(iii), the disintegrations 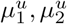 are Dirac almost everywhere, so each *π*_*u*_ reduces to a dirac mass matching the unique points of rank *u* in 𝒮_1_ and 𝒮_2_.

For this plan we have

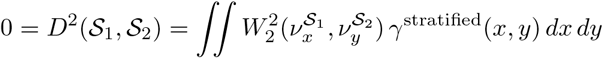

It is an upper bound of the second lower bound of Gromov–Wasserstein, minimized over all possible couplings:

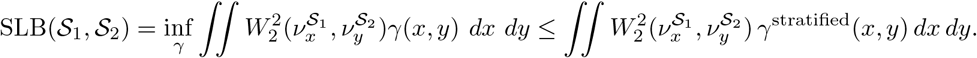

Thus the second lower bound of Gromov–Wasserstein distance is 0, and by its injectivity property [32], 𝒮_1_ and 𝒮_2_ must be isometric.

## Appendix B

Properties on stratified Wasserstein kernels

A Gaussian kernel defined by

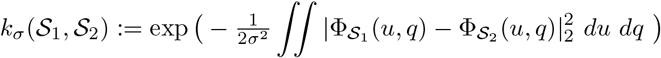

is positive semidefinite for all *σ >* 0 by the standard result that the Gaussian of a Hilbertian metric is PSD. Since Φ maps into a Euclidean space, ∥ · ∥_2_ is Hilbertian, hence *k*_*σ*_ is PSD.

If the embedding Φ is injective (cf. previous section), then *k*_*σ*_ is characteristic, i.e., it metrizes weak convergence of probability measures over the shape space when shapes are sampled i.i.d. from a distribution. Similarly, this extends to other kernels that are characteristic in Euclidean space, including Laplacian kernel and Matern kernels [43].

## Appendix C

Sample-based approximation is consistent

### Hypothesis (H2) (Bin growth)

*With n samples per shape, each divided into K equal-mass bins according to r*, min_*k*_ |*B*_*i,k*_| → ∞ *for i* = 1, 2 *so each bin gets a sufficiently large number of points*.

#### Theorem 3

**(Consistency of the stratified distance)** *Assume the existence and regularity of an oracle ranking functional (H1) and Bin Growth (H2). Let* 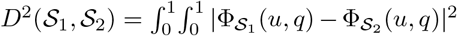 *denote the popu-lation distance*, 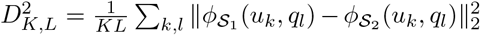, *and let* 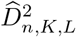 *be its empirical approximation with K equal-mass u-bins and an L-point q-grid computed when each shape has only n points observed*.

1. ***Fixed discretization***. *For any fixed* 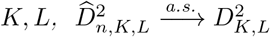 *as n* → ∞, *where* 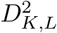 *is the corresponding discretized population functional*.
2. ***Refining discretization***. *As grid size* 1*/L* = max_*ℓ*_ (*q*_*ℓ*+1_ − *q*_*ℓ*_) → 0 *(refining q) and K* → ∞ *(refining u)*,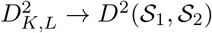.
3. ***Joint regime***. *If K* = *K*(*n*) → ∞ *and L* = *L*(*n*) → ∞ *with n/K*(*n*) = min_*k*_ |*B*_*i,k*_| → ∞, *then*

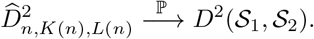

**Remark 3.1** *Part (a) ensures that with fixed small K, L the estimator is already consistent for its discretized target; parts (b)–(c) show that refining K, L recovers the continuous population distance*.

### Proof of consistency

Under Hypothesis (H1) and (H2), we aim to prove (a)-(c) as stated above.

#### Step 1

*Fixed K, L*. Let *U* = *F* (*r*) be the normalized rank. Equal-mass empirical bins *B*_*i,k*_ are order-statistic blocks in *U* ; by Rank/Bins and Bin Growth, *n*_*i,k*_ → ∞ and the empirical conditional law in *B*_*i,k*_ converges to the population conditional 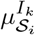. For each fixed *q*_*ℓ*_, the atomless, bounded-support assumption implies 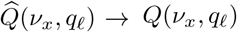 for every *x* (Glivenko–Cantelli Theorem). By the strong law within each bin,

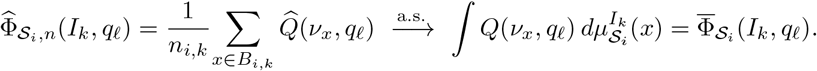

Since there are finitely many (*k, ℓ*), we obtain 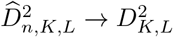.

#### Step 2

*Refining q*. For each fixed bin *I*_*k*_, the map 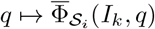 is bounded and monotone (average of quantile functions). Hence, as the mesh of the *q*-grid goes to zero, the Riemann sums converge to the integral:

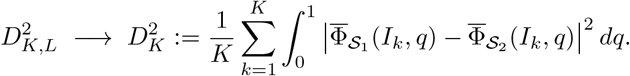

#### Step 3

*Refining u*.Let *G*(*u, q*) := Φ_*S*_ (*u, q*) − Φ_*S*_ (*u, q*) ∈ *L*^2^([0, 1]^2^). For each *q*, the bin-average 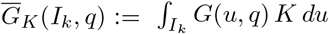 is the conditional expectation 𝔼 [*G*(*U, q*) | *σ*({*I*_*k*_})]. By the martingale convergence theorem for refining partitions, 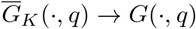 in *L*^2^([0, 1]) as *K* → ∞. Integrating in *q* and using dominated convergence yields 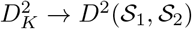.

#### Step 4

*Joint regime*. For *K* = *K*(*n*) and *L* = *L*(*n*), decompose

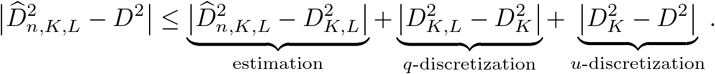

The estimation term → 0 by Step 1 applied with growing bin sizes (Hypothesis (H2)) and law of large numbers. The *q*- and *u*-discretization terms vanish by Steps 2 and 3 when the *L* → ∞ and *K* → ∞. Therefore 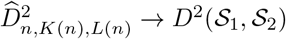.

## Appendix D

Computational cost

The oracle ranking function *r* is unknown in general. In experiments we approximate *r* via a finite-feature surrogate (e.g., a lexicographic combination of moments or quantiles). For any finite sample this map is injective for all but finitely many hyperparameter values (tiny *ϵ*), and its empirical bins converge to the population slices as sample size grows.

Let *N* be the number of sample points per shape, *M* the number of moments, *K* the number of bins, and *L* the number of discretization points for quantiles.

- **Local distance distributions:** Computing all *N* (*N* − 1)*/*2 pairwise distances is 𝒪 (*N* ^2^) in general; in low-dimensional ambient spaces this can be reduced using spatial data structures.
- **Moment computation:** With pairwise distances available, computing *M* moments for each point is 𝒪 (*MN*).
- **Quantile preparation:** To evaluate *L* quantiles for each *ν*_*x*_, one needs to sort each point’s *N* −1 distances in 𝒪 (*N* log *N*), giving 𝒪 (*N* ^2^ log *N*) total.
- **Ranking and binning:** Sorting *N* points by the ranking function *r*_*S*_(*x, ϵ*) is 𝒪 (*N* log *N*).
- **Bin-wise aggregation:** Sorting all distances within each bin (so *N* ^2^*/K* values) takes 𝒪 (*N* ^2^*/K* log(*N*)), then we can obtain the *L* quantiles.
- **Distance between shapes:** Once each shape is embedded as 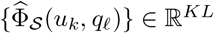, comparing two shapes takes 𝒪 (*KL*).

In practice, the quantile preparation and bin-wise aggregation can both be done through just a global sorting of all distances with cost 𝒪 (*N* ^2^ log *N*). If all steps are included, the dominant term is 𝒪 (*N* ^2^ log *N*) for exact quantiles, still much cheaper than a full Gromov–Wasserstein solve (𝒪 (*N* ^3^) or higher), while retaining injectivity under mild assumptions.

## Appendix E

Examples

This section has simple, 2D examples to help clarify how stratified distance works.

**Figure S1:**
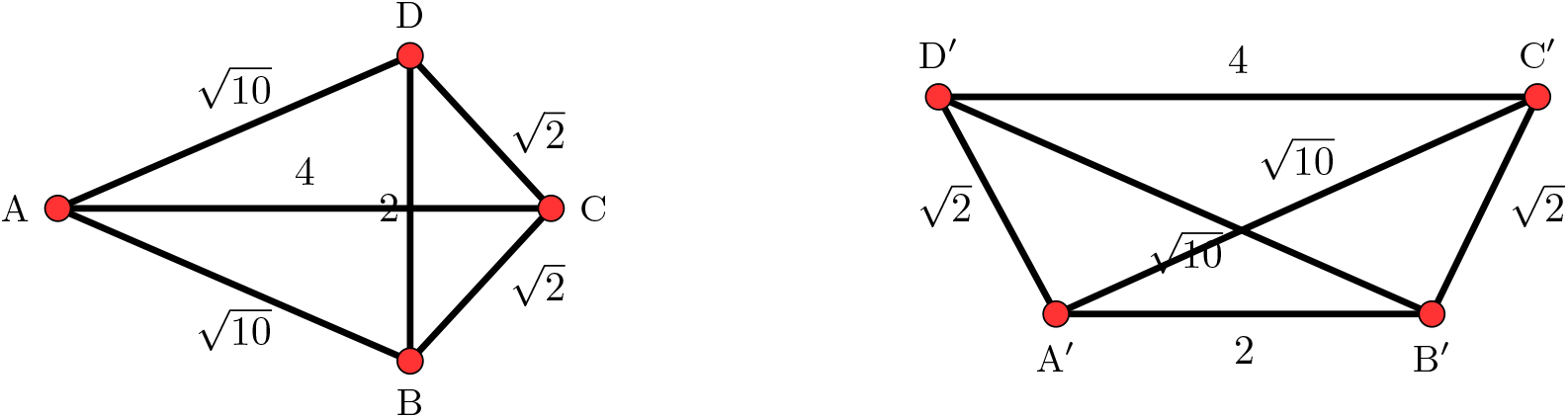
Illustration on global and local distance metrics.

### Appendix E.1 Difference between Wasserstein on global vs local distributions

Here we use a simple example taken from [6, 31] to illustrate the difference between local and global distance profiles:

The two shapes have distance matrices as below:

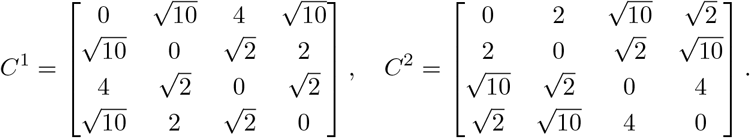

#### Global distances

thinking about global distances, both are uniformly distributed over 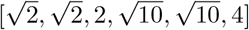, so the Wasserstein distance between global distances vanishes between these two shapes, indicating that the global Wasserstein distance cannot discriminate all shape spaces up to isometry.

#### Local distances

As a first step, the points should be ranked to create a correspondence. Sorted by average distance to all other points (descending order), the first shape results in *A, C, B, D* (with *B* and *D* interchangeable). The second shape is ordered by *C*^*′*^, *D*^*′*^, *A*^*′*^, *B*^*′*^, with *C*^*′*^ and *D*^*′*^ interchangeable and *A*^*′*^ and *B*^*′*^ interchangeable. As the local distances between *A* and *C*^*′*^ are already non-identical, the distance is nonzero.

### Appendix E.2 An example when sorting by mean can fail

In Figure S2, we show a five-point T-shape. For *a* ≈ 2.2666 the mean distance from the center *C* to the other points equals that from the right-arm point *A*, even though their local distance distributions differ. Including the second order information (variance) can help to tell that *A* is more likely to be on the ‘extreme’ side than *C* from its high variance.

**Figure S2:**
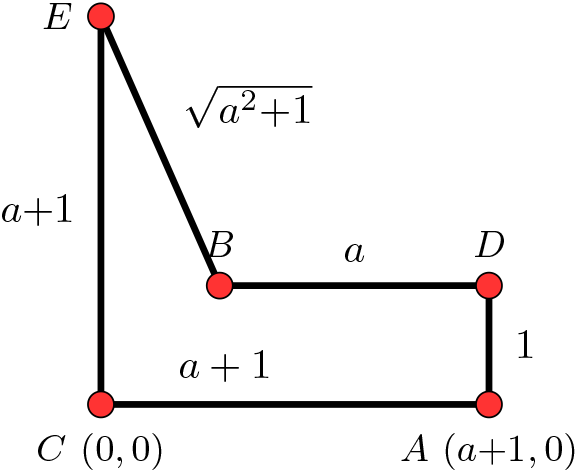
An example illustrating the effect of using information beyond the mean of local distance distributions.

### Appendix E.3 A comparison between all distances on random polygons

The second lower bound as well as the global Wasserstein distance over pairwise distances are both known to be a lower bound of Gromov–Wasserstein distance. The proposed stratified Wasserstein distance is an upper bound of the second lower bound, per the proof of injectivity. In this example, we perform a systematic comparison between these choices of distances on 200 random polygons with 6 sides, where the vertices are randomly sampled within the unit square. For the 200×200 pairs, we compute the distances between each pair of them, and visualize how the distances compare against each other in Figure S3.

As expected, the global Wasserstein distance and second lower bounds (SLB) do not exceed Gromov–Wasserstein distance, while the second lower bound offers greater power of discriminating the shape differences and have typically higher values compared to the global Wasserstein. This is consistent with the remark made by [32] that the second lower bound provides an upper bound of the global distance.

The stratified Wasserstein provides an upper bound on the lower bound of Gromov–Wasserstein distance, so in principle, there is no definite conclusion on the relationship between them. We remark that the stratified Wasserstein mostly serves as a lower bound of Gromov–Wasserstein, with very few rare cases of exceeding Gromov–Wasserstein, as shown in the middle panel of Figure S3. This agrees with the test power reported in the 2D cell contour example in Figure 2, panel (f) that stratified Wasserstein distance has slightly weaker test power.

**Figure S3:**
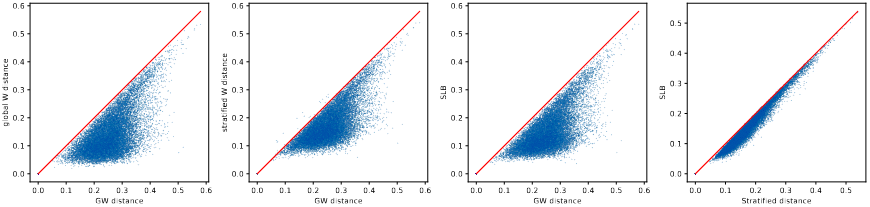
Comparison of distances across 200 × 200 pairs of random 6-sided polygons.

### Appendix E.4 Effect of imperfect matching by stratified functional

**Figure S4:**
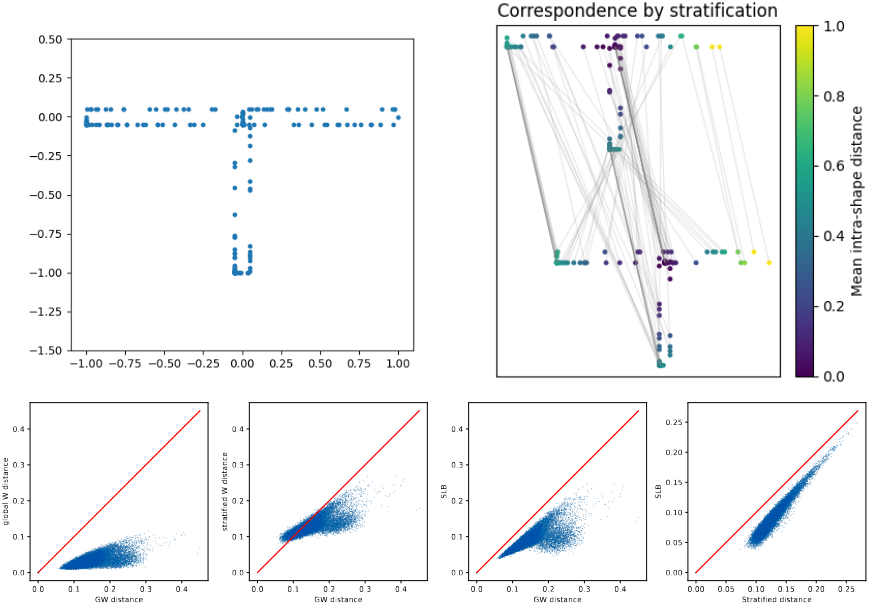
Top: T -shapes and example on false coupling by ranking the mean distance. Bottom: comparison across distances between 200 × 200 pairs of T -shapes with 100 randomly sampled points each.

Next, we manufacture an example where the incorrect matching for points can result in large deviations from the Gromov–Wasserstein distance and inflate the distance between shapes even if they are similar. Consider T-shapes with three equal arms, due to the symmetry, ranking by the mean distance will not yield the globally optimal coupling, as shown in the top right panel of Figure S4. As a result, the stratified distance is significantly greater than the second lower bound, and can still have relatively higher values even if the Gromov–Wasserstein distance is low. This suggests that stratified Wasserstein can over-penalize on the discrepancy if the functional (such as mean distance) does not yield appropriate correspondence between shapes, though such data with symmetry rarely seen in large-scale biological applications.

## Appendix F

Supplementary figures on experiments

**Figure S5:**
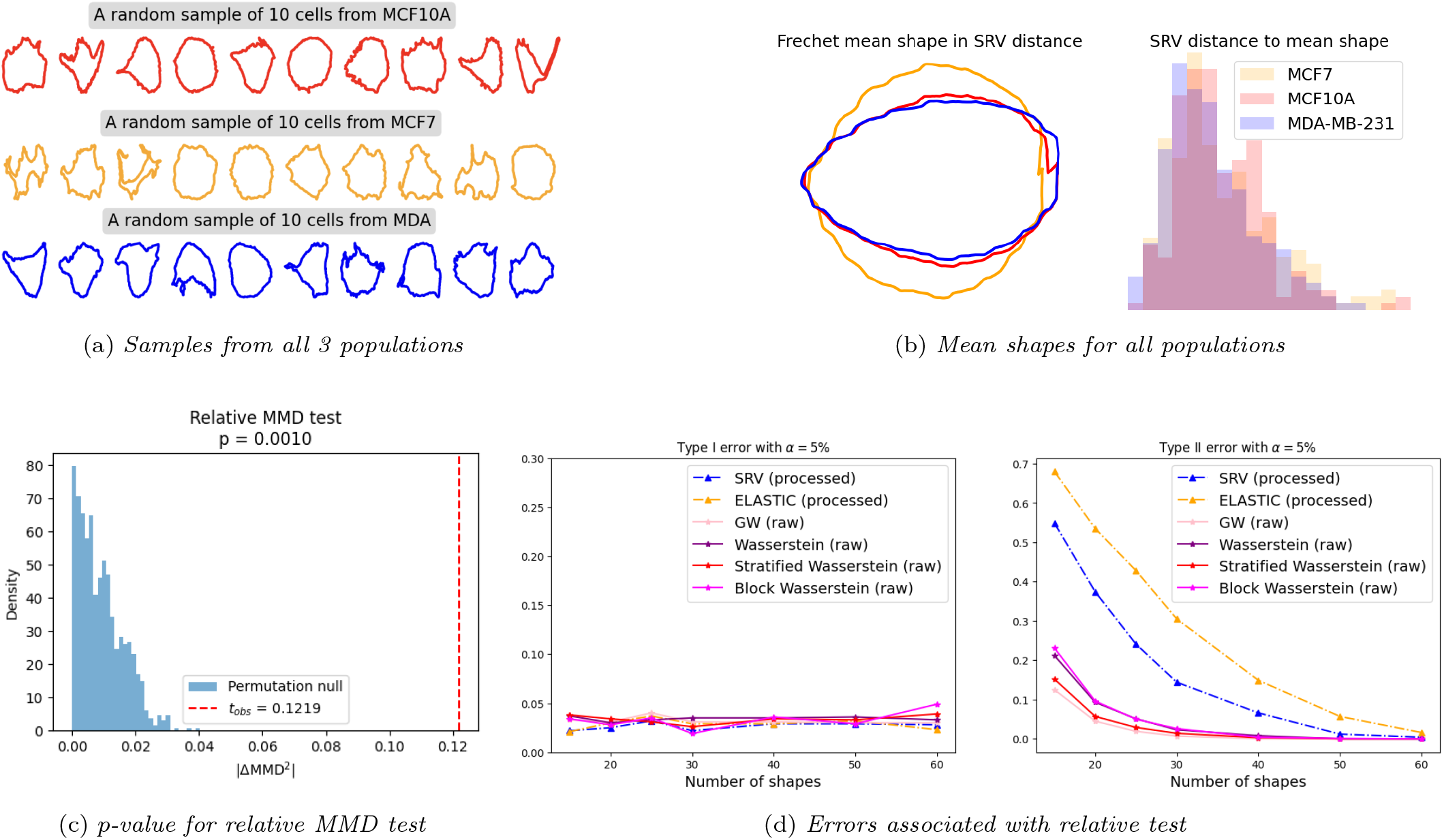
Supplementary results for all 3 populations in 2D cancer cell contour datasets. (a) Random samples from MCF10A (red), MCF7 (yellow), and MDA (blue). (b) Mean shapes for each category. (c) test statistic for whether cancer cell populations are closer than cancer versus normal. (d) Type I and II errors for different kernels for the same hypothesis testing.

**Figure S6:**
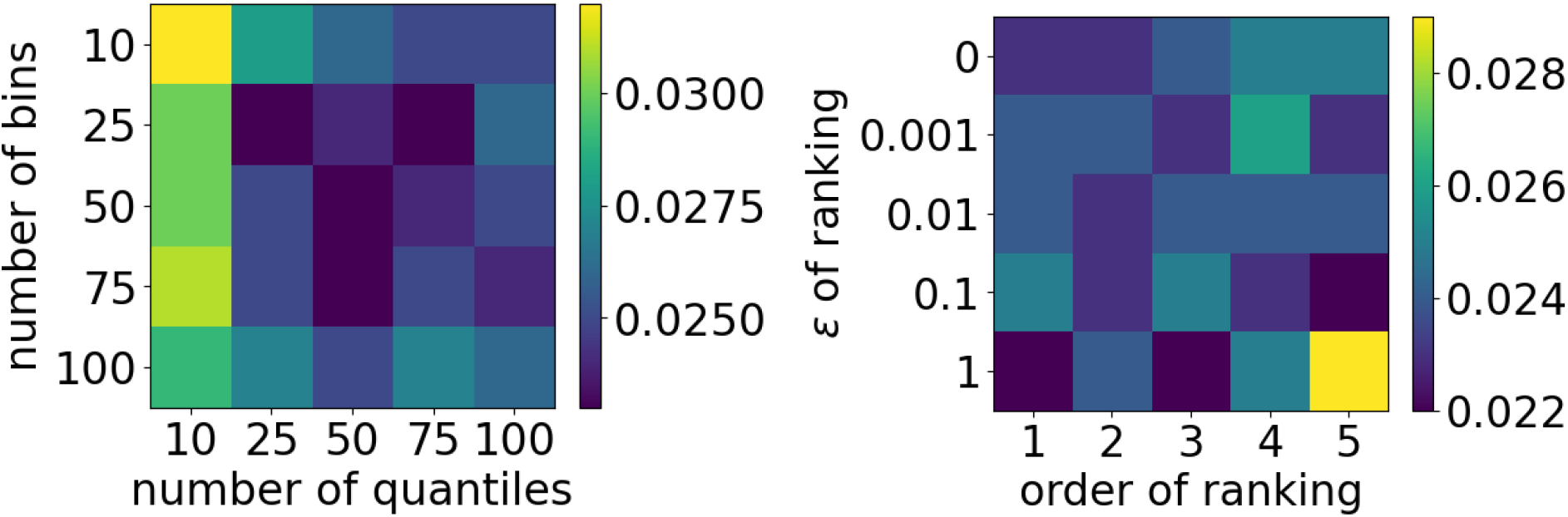
Type II error with 25 cancer cell shapes in 2D when varying hyperparameters, randomized over 1000 trials. Lower (Darker) is better. Left: varied discretization. Right: varied ranking functional. The results are robust to the choices of hyperparameters when the discretization is refined enough, and lower order of moments is better at respecting geometry and achieves lower error. We remark that the variation is in fact very small – as a reference, the Type II error of using SRV distance kernel is around 0.15.

**Figure S7:**
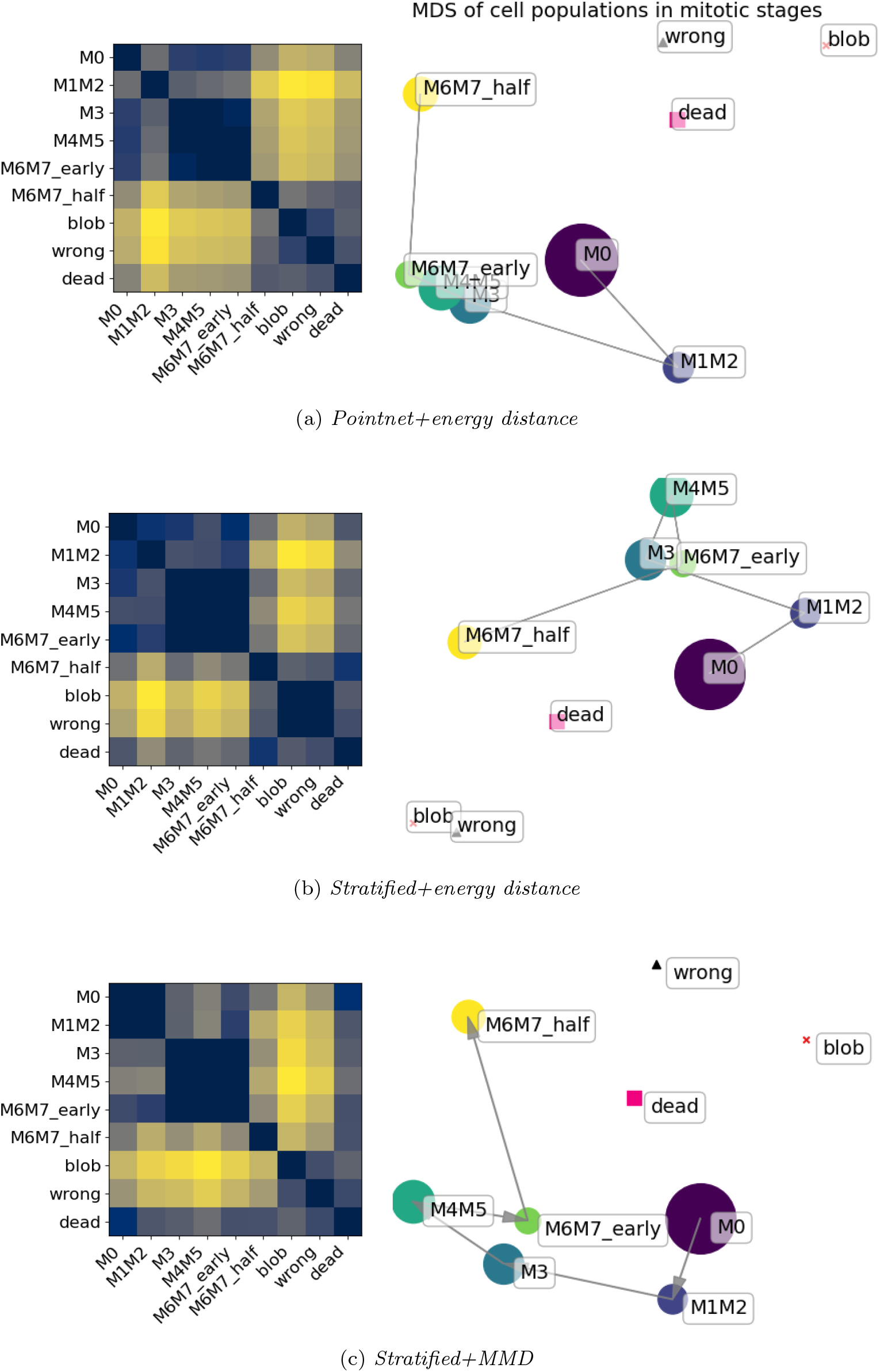
Comparison on population-level distances for mitotic dataset. Our framework shows a block-diagonal structure that sees similarity between consecutive stages, while both pointnet and stratified energy distance confuses M0 with intermediate stages such as M3 to M6M7 early.

## References

[1] E. Alizadeh, J. Castle, A. Quirk, C. D. Taylor, W. Xu, and A. Prasad. Cellular morphological features are predictive markers of cancer cell state. Computers in Biology and Medicine, 126:104044, 2020.

[2] E. Alizadeh, W. Xu, J. Castle, J. Foss, and A. Prasad. TISMorph: A tool to quantify texture, irregularity and spreading of single cells. PLoS One, 14(6):e0217346, June 2019.

[3] J. Altschuler, J. Weed, and P. Rigollet. Near-linear time approximation algorithms for optimal transport via sinkhorn iteration. In Proceedings of the 31st International Conference on Neural Information Processing Systems, NIPS’17, page 1961–1971, Red Hook, NY, USA, 2017. Curran Associates Inc.

[4] M. A. Astore, G. Woollard, D. Silva-Sánchez, W. Zhou, M. Kopylov, K. Dao Duc, R. R. Lederman, Y. Li, Y. Zhou, J. Yuan, F. Ye, Q. Gu, R. Vuillemot, S. Jonic, L. Dang, S. J. Ludtke, H. Bridges, S. Liu, M. McLean, V. Peretroukhin, J. Schwab, E. R. Cruz-Chú, P. Schwander, M. A. Gilles, A. Singer, D. Herreros, J.-M. Carazo, C. O. Sorzano, J. R. Feathers, E. D. Zhong, N. Grigorieff, P. Cossio, and S. M. Hanson. The inaugural flatiron institute cryo-em conformational heterogeneity challenge. July 2025.

[5] M. Bauer, M. Bruveris, S. Marsland, and P. W. Michor. Constructing reparameterization invariant metrics on spaces of plane curves. Differential Geometry and its Applications, 34: 139–165, 2014.

[6] M. Boutin and G. Kemper. On reconstructing n-point configurations from the distribution of distances or areas. Advances in Applied Mathematics, 32(4):709–735, May 2004.

[7] D. M. Boyer, Y. Lipman, E. St. Clair, J. Puente, B. A. Patel, T. Funkhouser, J. Jernvall, and I. Daubechies. Algorithms to automatically quantify the geometric similarity of anatomical surfaces. Proceedings of the National Academy of Sciences, 108(45):18221–18226, Oct. 2011.

[8] J. Burgess, J. J. Nirschl, M.-C. Zanellati, A. Lozano, S. Cohen, and S. Yeung-Levy. Orientation-invariant autoencoders learn robust representations for shape profiling of cells and organelles. Nature Communications, 15(1), Feb. 2024.

[9] M. Catalano and H. Lavenant. Hierarchical integral probability metrics: a distance on random probability measures with low sample complexity. In Proceedings of the 41st International Conference on Machine Learning, ICML’24. JMLR.org, 2024.

[10] M. Cuturi. Sinkhorn distances: Lightspeed computation of optimal transport. In C. Burges, L. Bottou, M. Welling, Z. Ghahramani, and K. Weinberger, editors, Advances in Neural Information Processing Systems, volume 26. Curran Associates, Inc., 2013.

[11] K. Dao Duc, S. S. Batra, N. Bhattacharya, J. H. D. Cate, and Y. S. Song. Differences in the path to exit the ribosome across the three domains of life. Nucleic Acids Research, 47(8):4198–4210, Feb. 2019.

[12] I. L. Dryden and K. V. Mardia. Statistical Shape Analysis, with Applications in R. Wiley, Sept. 2016.

[13] P. Dvurechensky, A. Gasnikov, and A. Kroshnin. Computational optimal transport: Complexity by accelerated gradient descent is better than by sinkhorn’s algorithm. In J. Dy and A. Krause, editors, Proceedings of the 35th International Conference on Machine Learning, volume 80 of Proceedings of Machine Learning Research, pages 1367–1376. PMLR, 10–15 Jul 2018.

[14] M. Ekvall, L. Bergenstråhle, A. Andersson, P. Czarnewski, J. Olegård, L. Käll, and J. Lundeberg. Spatial landmark detection and tissue registration with deep learning. Nature Methods, 21(4):673–679, Mar. 2024.

[15] R. Flamary, N. Courty, A. Gramfort, M. Z. Alaya, A. Boisbunon, S. Chambon, L. Chapel, A. Corenflos, K. Fatras, N. Fournier, L. Gautheron, N. T. Gayraud, H. Janati, A. Rakotomamonjy, I. Redko, A. Rolet, A. Schutz, V. Seguy, D. J. Sutherland, R. Tavenard, A. Tong, and T. Vayer. Pot: Python optimal transport. Journal of Machine Learning Research, 22(78): 1–8, 2021.

[16] R. Flamary, C. Vincent-Cuaz, N. Courty, A. Gramfort, O. Kachaiev, H. Quang Tran, L. David, C. Bonet, N. Cassereau, T. Gnassounou, E. Tanguy, J. Delon, A. Collas, S. Mazelet, L. Chapel, T. Kerdoncuff, X. Yu, M. Feickert, P. Krzakala, T. Liu, and E. Fernandes Montesuma. Pot python optimal transport (version 0.9.5), 2024.

[17] N. Fournier and A. Guillin. On the rate of convergence in wasserstein distance of the empirical measure. Probability Theory and Related Fields, 162(3–4):707–738, Oct. 2014.

[18] M. Franchi, Z. Piperigkou, K.-A. Karamanos, L. Franchi, and V. Masola. Extracellular matrix-mediated breast cancer cells morphological alterations, invasiveness, and microvesicles/exosomes release. Cells, 9(9):2031, Sept. 2020.

[19] K. W. Govek, P. Nicodemus, Y. Lin, J. Crawford, A. B. Saturnino, H. Cui, K. Zoga, M. P. Hart, and P. G. Camara. Cajal enables analysis and integration of single-cell morphological data using metric geometry. Nature Communications, 14(1), June 2023.

[20] A. Gretton, K. M. Borgwardt, M. J. Rasch, B. Schölkopf, and A. Smola. A kernel two-sample test. J. Mach. Learn. Res., 13(null):723–773, Mar. 2012.

[21] A. Gretton, K. Fukumizu, C. Teo, L. Song, B. Schölkopf, and A. Smola. A kernel statistical test of independence. In J. Platt, D. Koller, Y. Singer, and S. Roweis, editors, Advances in Neural Information Processing Systems, volume 20. Curran Associates, Inc., 2007.

[22] M. Kazhdan, T. Funkhouser, and S. Rusinkiewicz. Rotation invariant spherical harmonic representation of 3d shape descriptors. In Proceedings of the 2003 Eurographics/ACM SIGGRAPH Symposium on Geometry Processing, SGP ‘03, page 156–164, Goslar, DEU, 2003. Eurographics Association.

[23] D. G. Kendall. Shape manifolds, procrustean metrics, and complex projective spaces. Bulletin of the London Mathematical Society, 16(2):81–121, Mar. 1984.

[24] S. Kolouri, Y. Zou, and G. K. Rohde. Sliced wasserstein kernels for probability distributions, 2015.

[25] N. Kravtsova. The np-hardness of the gromov-wasserstein distance, 2024.

[26] W. Li. breast cancer analysis.ipynb. GitHub, 2024. cited 2025-10-12; commit 92c7a58.

[27] W. Li, A. Prasad, N. Miolane, and K. Dao Duc. Unveiling cellular morphology: statistical analysis using a riemannian elastic metric in cancer cell image datasets. Information Geometry, 7(S2):845–859, Aug. 2024.

[28] Y.-S. Liu, Q. Li, G.-Q. Zheng, K. Ramani, and W. Benjamin. Using diffusion distances for flexible molecular shape comparison. BMC Bioinformatics, 11(1), Sept. 2010.

[29] N. Miolane, N. Guigui, A. L. Brigant, J. Mathe, B. Hou, Y. Thanwerdas, S. Heyder, O. Peltre, N. Koep, H. Zaatiti, H. Hajri, Y. Cabanes, T. Gerald, P. Chauchat, C. Shewmake, D. Brooks, B. Kainz, C. Donnat, S. Holmes, and X. Pennec. Geomstats: A python package for riemannian geometry in machine learning. Journal of Machine Learning Research, 21(223): 1–9, 2020.

[30] E. Munch. A user’s guide to topological data analysis. Journal of Learning Analytics, 4(2), July 2017.

[31] F. Mémoli. Gromov–wasserstein distances and the metric approach to object matching. Foundations of Computational Mathematics, 11(4):417–487, Apr. 2011.

[32] F. Mémoli and T. Needham. Distance distributions and inverse problems for metric measure spaces. Studies in Applied Mathematics, 149(4):943–1001, Aug. 2022.

[33] T. Needham and S. Kurtek. Simplifying transforms for general elastic metrics on the space of plane curves. SIAM Journal on Imaging Sciences, 13(1):445–473, Jan. 2020.

[34] R. M. Neve, K. Chin, J. Fridlyand, J. Yeh, F. L. Baehner, T. Fevr, L. Clark, N. Bayani, J.-P. Coppe, F. Tong, T. Speed, P. T. Spellman, S. DeVries, A. Lapuk, N. J. Wang, W.-L. Kuo, J. L. Stilwell, D. Pinkel, D. G. Albertson, F. M. Waldman, F. McCormick, R. B. Dickson, M. D. Johnson, M. Lippman, S. Ethier, A. Gazdar, and J. W. Gray. A collection of breast cancer cell lines for the study of functionally distinct cancer subtypes. Cancer Cell, 10(6):515–527, Dec. 2006.

[35] R. Osada, T. Funkhouser, B. Chazelle, and D. Dobkin. Shape distributions. ACM Transactions on Graphics, 21(4):807–832, Oct. 2002.

[36] R. C. Perez, S. da Veiga, J. Garnier, and B. Staber. Gaussian process regression with sliced wasserstein weisfeiler-lehman graph kernels, 2024.

[37] G. Peyré and M. Cuturi. Computational optimal transport. 2018.

[38] R. Pogodin, N. Deka, Y. Li, D. J. Sutherland, V. Veitch, and A. Gretton. Efficient conditionally invariant representation learning. In The Eleventh International Conference on Learning Representations, 2023.

[39] C. R. Qi, H. Su, K. Mo, and L. J. Guibas. Pointnet: Deep learning on point sets for 3d classification and segmentation. arXiv preprint arXiv:1612.00593, 2016.

[40] C. R. Qi, L. Yi, H. Su, and L. J. Guibas. Pointnet++: deep hierarchical feature learning on point sets in a metric space. In Proceedings of the 31st International Conference on Neural Information Processing Systems, NIPS’17, page 5105–5114, Red Hook, NY, USA, 2017. Curran Associates Inc.

[41] G. Rioux, Z. Goldfeld, and K. Kato. Entropic gromov-wasserstein distances: stability and algorithms. J. Mach. Learn. Res., 25(1), Jan. 2024.

[42] M. Scetbon, G. Peyré, and M. Cuturi. Linear-time gromov Wasserstein distances using low rank couplings and costs. In K. Chaudhuri, S. Jegelka, L. Song, C. Szepesvari, G. Niu, and S. Sabato, editors, Proceedings of the 39th International Conference on Machine Learning, volume 162 of Proceedings of Machine Learning Research, pages 19347–19365. PMLR, 17–23 Jul 2022.

[43] D. Sejdinovic, B. Sriperumbudur, A. Gretton, and K. Fukumizu. Equivalence of distance-based and rkhs-based statistics in hypothesis testing. The Annals of Statistics, 41(5), Oct. 2013.

[44] L. Song, A. Smola, A. Gretton, J. Bedo, and K. Borgwardt. Feature selection via dependence maximization. J. Mach. Learn. Res., 13(1):1393–1434, May 2012.

[45] B. K. Sriperumbudur, A. Gretton, K. Fukumizu, B. Schölkopf, and G. R. G. Lanckriet. Hilbert space embeddings and metrics on probability measures, 2009.

[46] A. Srivastava, E. Klassen, S. H. Joshi, and I. H. Jermyn. Shape analysis of elastic curves in euclidean spaces. IEEE Transactions on Pattern Analysis and Machine Intelligence, 33(7):1415–1428, July 2011.

[47] M. Styner, I. Oguz, S. Xu, C. Brechbühler, D. Pantazis, J. J. Levitt, M. E. Shenton, and G. Gerig. Framework for the statistical shape analysis of brain structures using SPHARM-PDM. Insight J., (1071):242–250, 2006.

[48] D. J. Sutherland and N. Deka. Unbiased estimators for the variance of mmd estimators, 2019.

[49] M. P. Viana, J. Chen, T. A. Knijnenburg, R. Vasan, C. Yan, J. E. Arakaki, M. Bailey, B. Berry, A. Boren-sztejn, E. M. Brown, S. Carlson, J. A. Cass, B. Chaudhuri, K. R. Cordes Metzler, M. E. Coston, Z. J. Crabtree, S. Davidson, C. M. DeLizo, S. Dhaka, S. Q. Dinh, T. P. Do, J. Domingus, R. M. Donovan-Maiye, A. J. Ferrante, T. J. Foster, C. L. Frick, G. Fujioka, M. A. Fuqua, J. L. Gehring, K. A. Gerbin, T. Gran-charova, B. W. Gregor, L. J. Harrylock, A. Haupt, M. C. Hendershott, C. Hookway, A. R. Horwitz, H. C. Hughes, E. J. Isaac, G. R. Johnson, B. Kim, A. N. Leonard, W. W. Leung, J. J. Lucas, S. A. Ludmann, A. M. Lyons, H. Malik, R. McGregor, G. E. Medrash, S. L. Meharry, K. Mitcham, I. A. Mueller, T. L. Murphy-Stevens, A. Nath, A. M. Nelson, S. A. Oluoch, L. Paleologu, T. A. Popiel, M. M. Riel-Mehan, B. Roberts, L. M. Schaefbauer, M. Schwarzl, J. Sherman, S. Slaton, M. F. Sluzewski, J. E. Smith, Y. Sul, M. J. Swain-Bowden, W. J. Tang, D. J. Thirstrup, D. M. Toloudis, A. P. Tucker, V. Valencia, W. Wiegraebe, T. Wijeratna, R. Yang, R. J. Zaunbrecher, R. L. D. Labitigan, A. L. Sanborn, G. T. Johnson, R. N. Gunawardane, N. Gaudreault, J. A. Theriot, and S. M. Rafelski. Integrated intracellular organization and its variations in human ips cells. Nature, 613(7943):345–354, Jan. 2023.

[50] C. E. Walczak, S. Cai, and A. Khodjakov. Mechanisms of chromosome behaviour during mitosis. Nature Reviews Molecular Cell Biology, 11(2):91–102, Jan. 2010.

[51] K. Zhang, J. Peters, D. Janzing, and B. Schölkopf. Kernel-based conditional independence test and application in causal discovery. In Proceedings of the Twenty-Seventh Conference on Uncertainty in Artificial Intelligence, UAI’11, page 804–813, Arlington, Virginia, USA, 2011. AUAI Press.

[52] R. Zhang, R. T. Ogden, M. Picard, and A. Srivastava. Nonparametric k-sample test on shape spaces with applications to mitochondrial shape analysis. Journal of the Royal Statistical Society Series C: Applied Statistics, 71(1):51–69, Jan. 2022.

